# ACKR3–arrestin2/3 complexes reveal molecular consequences of GRK-dependent barcoding

**DOI:** 10.1101/2023.07.18.549504

**Authors:** Qiuyan Chen, Christopher T. Schafer, Somnath Mukherjee, Martin Gustavsson, Parth Agrawal, Xin-Qiu Yao, Anthony A. Kossiakoff, Tracy M. Handel, John J. G. Tesmer

## Abstract

Atypical chemokine receptor 3 (ACKR3, also known as CXCR7) is a scavenger receptor that regulates extracellular levels of the chemokine CXCL12 to maintain responsiveness of its partner, the G protein-coupled receptor (GPCR), CXCR4. ACKR3 is notable because it does not couple to G proteins and instead is completely biased towards arrestins. Our previous studies revealed that GRK2 and GRK5 install distinct distributions of phosphates (or “barcodes”) on the ACKR3 carboxy terminal tail, but how these unique barcodes drive different cellular outcomes is not understood. It is also not known if arrestin2 (Arr2) and 3 (Arr3) bind to these barcodes in distinct ways. Here we report cryo-electron microscopy structures of Arr2 and Arr3 in complex with ACKR3 phosphorylated by either GRK2 or GRK5. Unexpectedly, the finger loops of Arr2 and 3 directly insert into the detergent/membrane instead of the transmembrane core of ACKR3, in contrast to previously reported “core” GPCR–arrestin complexes. The distance between the phosphorylation barcode and the receptor transmembrane core regulates the interaction mode of arrestin, alternating between a tighter complex for GRK5 sites and heterogenous primarily “tail only” complexes for GRK2 sites. Arr2 and 3 bind at different angles relative to the core of ACKR3, likely due to differences in membrane/micelle anchoring at their C-edge loops. Our structural investigations were facilitated by Fab7, a novel Fab that binds both Arr2 and 3 in their activated states irrespective of receptor or phosphorylation status, rendering it a potentially useful tool to aid structure determination of any native GPCR–arrestin complex. The structures provide unprecedented insight into how different phosphorylation barcodes and arrestin isoforms can globally affect the configuration of receptor–arrestin complexes. These differences may promote unique downstream intracellular interactions and cellular responses. Our structures also suggest that the 100% bias of ACKR3 for arrestins is driven by the ability of arrestins, but not G proteins, to bind GRK-phosphorylated ACKR3 even when excluded from the receptor cytoplasmic binding pocket.

## INTRODUCTION

Chemokines control the migration and localization of leukocytes and play fundamental roles in regulating immune and inflammatory responses including inflammation associated with cancer. One such chemokine, CXCL12, functions by binding to two 7 transmembrane domain (7TM) receptors: C-X-C chemokine receptor type 4 (CXCR4), and atypical chemokine receptor 3 (ACKR3). CXCR4 activates heterotrimeric Gα_i_ proteins and regulates cell movement. It is the subject of numerous clinical trials for leukemia, lymphoma, and solid tumors (clinicaltrials.gov) as it promotes multiple steps in the growth of primary tumors and progression to metastatic disease (Balkwill, 2004; Burns et al., 2006; Miao et al., 2007; Sanchez-Martin et al., 2013). Like CXCR4, ACKR3 is upregulated in many cancers, as well as on endothelial cells in the tumor vasculature where it cooperates with CXCR4 to promote the cancer phenotype (Balkwill, 2004). However in most cells, ACKR3 does not activate G proteins (Meyrath et al., 2020; Naumann et al., 2010; Odemis et al., 2010), but nevertheless is phosphorylated by GRKs and robustly recruits arrestins in response to CXCL12 (Canals et al., 2012; Levoye et al., 2009; Rajagopal et al., 2010; Torossian et al., 2014). One of its primary roles is to scavenge chemokine by internalizing with CXCL12, trafficking the chemokine to lysosomes for degradation and recycling back to the membrane for further rounds of ligand consumption (Naumann et al., 2010; Thelen and Thelen, 2008). This process regulates the levels of extracellular chemokine and is crucial for maintaining CXCR4 responsiveness in the context of normal physiology (Saaber et al., 2019) and tumor metastasis (Luker et al., 2012). ACKR3 has also been shown to physically interact with Connexin 43 and regulate the gap junction protein in an arrestin-dependent manner. These interactions may have broad implications for the role of ACKR3 in many processes particularly in the brain, including brain cancers (Fumagalli et al., 2020; Neves et al., 2019).

The principal objective of this study was to understand the bias of ACKR3 for arrestins and the molecular basis of arrestin activation by CXCL12 through ACKR3. We previously showed that GRK2 and GRK5, kinases representing the two major human subfamilies, phosphorylate ACKR3 in an activation-dependent fashion at different regions of its cytoplasmic tail (**Figure 1A**), giving rise to distinct phosphorylation “barcodes”. We also showed that GRK2-mediated phosphorylation only occurs with coactivation of CXCR4 whereas GRK5 phosphorylation is operative when ACKR3 is expressed alone, suggesting functional consequences of different barcodes (Schafer et al., 2023). The barcodes could also differentially regulate chemokine scavenging by ACKR3, which is dependent on phosphorylation of the receptor C-terminus (Hoffmann et al., 2012; Saaber et al., 2019). Finally, we found that phosphorylation of ACKR3 by either kinase enhances its binding not only to arrestin3 (Arr3), but also to arrestin2 (Arr2), setting the stage for comparison of the molecular consequences of GRK barcoding vs. arrestin isoform at the same receptor. Towards this end, our study provides evidence of large differences in the configuration of ACKR3– arrestin complexes that result from not only different GRK isoform barcoding but also the two arrestin isoforms. These distinct configurations could be selectively targeted to block specific functions of ACKR3.

**Figure 1.**
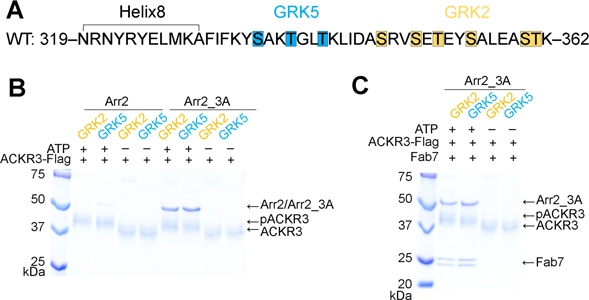
Arrestin binding to ACKR3 depends on GRK2/5 phosphorylation. (A) Sequence of the ACKR3 C-tail with the observed GRK2 phosphorylation sites highlighted in orange and the *unique* sites observed for GRK5 in blue. (B) A flag pulldown assay shows that comparable amounts of Arr2_3A, but not WT Arr2, binds to ACKR3 phosphorylated by GRK2 or GRK5. The interaction is abolished when ACKR3 is not phosphorylated. (C) Flag pulldown assays show that Fab7 coelutes with pACKR3(GRK2)–Arr2_3A and pACKR3(GRK5)–Arr2_3A. This interaction is dependent on GRK activity because non-phosphorylated ACKR3 does not pulldown Arr2_3A or Fab7.

## RESULTS AND DISCUSSION

### GRK Phosphorylation Is Required for Arrestin Coupling to ACKR3

Both GRK2 in the presence of G_ýy_, and GRK5 in the presence of PIP_2_, efficiently phosphorylate ACKR3 but install phosphates in distinct regions of the ACKR3 C-tail (**Figure 1A**) (Schafer et al., 2023). GRK2 phosphorylates Ser and Thr residues more distal to the transmembrane (TM) core, consistent with the results of (Zarca et al., 2021), whereas GRK5 installs phosphates beginning at more proximal sites, and under some conditions, sites that overlap with those of GRK2 (Schafer et al., 2023). Thus, we reasoned that this system would allow us to assess whether distinct phosphorylation patterns installed by the two GRK isoforms confer distinct structural responses in the two major arrestin isoforms, Arr2 and Arr3, that could lead to distinct downstream outcomes. To this end, we first assessed how phosphorylation affects arrestin binding using pulldown assays (**Figure 1B**). We tested both wildtype (WT) and preactivated forms of Arr2 and 3: Arr2_3A (I386A, V387A, F388A mutations in the C-tail), and Arr3_ΔC (truncated after residue 392) (Gurevich, 1998), henceforth referred to as Arr2 and Arr3 unless otherwise noted. Comparable amounts of Arr2 and Arr3 were pulled down with GRK2 or GRK5 phosphorylated ACKR3 (**Figure 1B**). Unphosphorylated ACKR3 failed to pull-down any detectable amount of Arr2 or Arr3 (**Figure 1B**), indicating that GRK phosphorylation, regardless of isoform, is required for efficient arrestin recruitment to activated ACKR3 (Schafer et al., 2023).

### A Novel Fab7 Facilitates cryo-EM Studies of pACKR3-Arr2/3 Complexes

Although complexes of phosphorylated ACKR3 (pACKR3) in complex with Arr2/3 could be isolated for cryo-EM studies, 2D class averages showed poorly defined features consistent with conformational heterogeneity. This was not surprising because arrestins gain substantial conformational flexibility upon full activation (Zhuang et al., 2013). To stabilize the complexes and assist in structure determinations, we performed a phage display selection to screen for novel Fabs that selectively bind to arrestin in its activated state regardless of an activating receptor peptide. For this purpose, we used pre-activated Arr3_ΔC in complex with the small molecule activator inositol hexakisphosphate (IP_6_). The IP_6_ binding site overlaps with the receptor phosphate binding sites (Chen et al., 2017). Moreover, IP_6_ promotes trimerization of Arr3 via its receptor-binding interfaces, which decreases the possibility of selecting Fabs that directly compete with receptor binding. Of our lead Fabs, we found that one, Fab7, binds to Arr3 as well as Arr2 while in complex with ACKR3 whether it is phosphorylated by either GRK2 or GRK5 (**Figure 1C**).

### Structure Determinations of GRK2/5 Phosphorylated ACKR3 in Complex with Arr2/3

We then determined cryo-EM structures for a series of ACKR3−arrestin complexes (**Figure 2A, B, Table S1**). In all structures, ACKR3 is bound to the chemokine mutant CXCL12_LRHQ_, a variant with a four residue substitution at the N-terminus (Leu-Arg-His-Gln) that prolongs its residence time on the receptor relative to WT CXCL12 (Gustavsson et al., 2019; Hanes et al., 2015). The main comparative set of structures include GRK5 phosphorylated ACKR3 (pACKR3(GRK5)) and GRK2 phosphorylated ACKR3 (pACKR3(GRK2)) in complex with either Arr2 or Arr3, all in LMNG/CHS micelles (**Figure 2A, S1-4, Table S1**). In parallel, we determined the structure of Arr2 in complex with pACKR3(GRK2) in POPC/POPS nanodiscs (**Figure 2B, S5**), wherein Fab7 was further stabilized with an anti-Fab nanobody (Nb) that binds to the elbow of the Fab light chain (Ereno-Orbea et al., 2018). In all five reconstructions, Fab7 binds to the interdomain hinge region of Arr2 and 3, opposite to the finger loop (**Figure 2A, B**). In none of our structures does Fab7 interact with the phosphorylated tail or TM domain of ACKR3, suggesting that it would interact with GPCR–arrestin complexes irrespective of GPCR identity, barcoding, and arrestin isoform. In contrast, Fab30, a Fab used to stabilize most other reported GPCR–arrestin structures, interacts with the hinge region of arrestin and a phosphorylated residue in the bound receptor tail (Shukla et al., 2013). Fab30 fails to bind to arrestins bound to phosphopeptides derived from the ACKR3 C-tail (Maharana et al., 2023b; Sarma et al., 2022).

**Figure 2.**
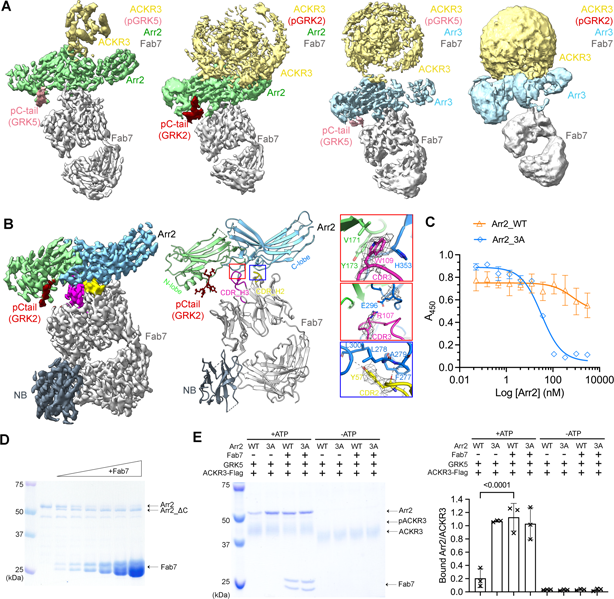
Structural and functional characterization of Fab7, a new arrestin conformational sensor. (A) Sharpened maps of the 3.2 Å pACKR3(GRK5)–Arr2–Fab7, the 3.5 Å pACKR3(GRK2)–Arr2–Fab7, the 3.9 Å pACKR3(GRK5)–Arr3–Fab7, and the 7.3 Å pACKR3(GRK2)–Arr3–Fab7 complexes. ACKR3 in all four complexes is solubilized in LMNG/CHS detergent micelles. (B) Sharpened map and model of the 3.0 Å pACKR3(GRK2)–Arr2– Fab7–NB in POPC/POPS nanodiscs (PDB entry XXXX). The density of the ACKR3 TM domain and the nanodisc is not evident. Insets show interfacial details discussed in the text. (C) ELISA analysis of Fab7 competition assay reveals that preactivated Arr2 (IC50 ∼35 nM) competes for Fab7 binding more efficiently than Arr2 WT (IC50 > 5 µM). Error bars represent S.D. from three technical replicates. (D) Limited trypsin digestion of Arr2 WT in the presence of increasing concentrations of Fab7. (E) A Flag pulldown assay shows that Fab7 significantly promotes Arr2 WT binding to pACKR3(GRK5), but it does not increase Arr2 WT or 3A binding non-phosphorylated ACKR3. One representative gel is shown. The ratios between the density of bound Arr2 and that of bound ACKR3 are compared using one-way ANOVA followed by a Dunnett’s multiple comparison test (P < 0.0001). Error bars represent S.D. from three technical replicates.

### Fab7 Interface

ACKR3 was not evident in the pACKR3(GRK2)–Arr2 nanodisc structure except for its phosphopeptide region (residues 351-358; **Figure 2B**), but the map for Arr2–Fab7–Nb was the highest resolution among our models (3 Å, **Figure S5, Table S1**) and allowed us to accurately map the structure of Fab7 and its interface with Arr2. Fab7 binds entirely with its heavy chain using all three complementarity-determining regions (CDRs), although the light chain CDR-L3 is indirectly involved. The most prominent interactions are formed by CDR-H3, which contains an 18-residue loop that binds in the hinge of Arr2 (**Figure 2B**). Fab7-Trp109 interacts with His353 on the C-lobe and Val171 and Tyr173 on the N-lobe of Arr2, while Fab7-Arg107 forms a salt bridge with Glu296 on the C-lobe. Another cluster of contacts outside the hinge is provided by Fab7-Tyr57 in CDR-H2, which forms nonpolar contacts with Arr2-Phe277 and - Leu278, and hydrogen bonds with the backbone nitrogens of Arr2-Ala279 and - Leu300 (**Figure 2B**). These same interactions are also formed by Fab30-Tyr57, which has an identical CDR2. However, Fab7 has less extensive interaction with Arr2 (buried surface area ∼900 Å^2^) than Fab30 (buried surface area ∼700 Å^2^).

The Fab7 interface is not compatible with the basal conformation of Arr2 in part because Fab7-Tyr57 and -Trp109 would be excluded via steric clashes in their respective pockets (**Figure S6A**). This suggests that Fab7 should strongly favor the activated state of arrestin. To test this experimentally, we performed a competitive multipoint protein ELISA where the binding of Fab7 to Arr3_ι1C·IP_6_ was competed with increasing concentrations of WT Arr2 or Arr2_3A (**Figure 2C**). Arr2_3A displaced Fab7 with an IC_50_ of ∼35 nM, whereas WT Arr2 barely competed with an IC_50_ > 5 µM. Similar results were obtained for WT Arr3 vs. Arr3_ι1C (**Figure S6B**). To check whether Fab7 can activate Arr2 on its own, we assessed how Fab7 binding affects trypsin digestion of Arr2. Fab7 facilitated digestion, producing C-terminally truncated Arr2 (**Figure 2D**). This suggests that Fab7 traps Arr2 when its C-tail is dissociated, rendering it more susceptible to proteolysis. Finally, we tested whether Fab7 boosts Arr2 binding to ACKR3 via pulldown assays (**Figure 2E**). Fab7 significantly increased the binding of WT Arr2 to phosphorylated ACKR3; however, it did not promote the binding of either WT or preactivated Arr2 to unphosphorylated ACKR3 (**Figure 2E**). Taken together, we conclude that Fab7 selectively binds arrestins in their activated state but can only drive arrestin activation independent of ACKR3 at high saturating concentrations, as evidenced by proteolytic protection assays (**Figure 2D**).

### The pACKR3(GRK5)–Arr2 Complex Reveals a Novel Arrestin–Receptor Configuration

The pACKR3(GRK5)–Arr2 complex yielded the highest resolution micelle complex (3.2 Å) and the strongest receptor density (**Figure 2A, 3A, S1**). The cytoplasmic ends of the seven TM helices of ACKR3 are the most well resolved and resemble those in the open active state CID24–CXCL12_LRHQ_– ACKR3 complex (PDB entry 7SK6) (Yen et al., 2022). There is also weak density for intracellular loop 3 (ICL3), for the first three residues of CXCL12_LRHQ_ in the orthosteric pocket (although the entire chemokine was left in the model), and for some cholesterol molecules that were commonly observed in previously published structures of ACKR3 (**Figure 3A**) (Yen et al., 2022). Helix8 of the receptor is not ordered (**Figure 3A**).

**Figure 3.**
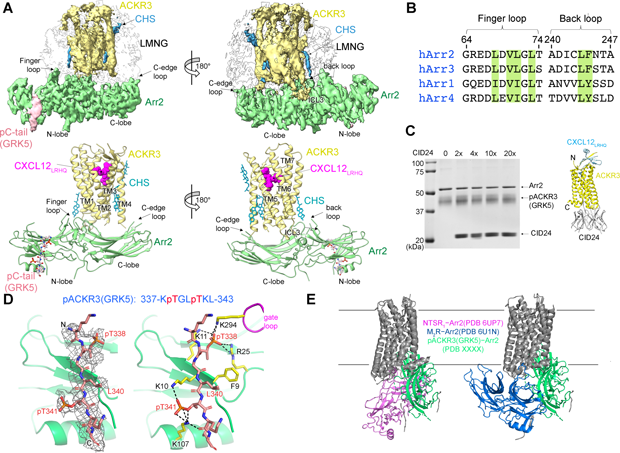
The interface between pACKR3(GRK5) and Arr2 is supported by a novel and several conventional interactions. (A) Sharpened maps and models of pACKR3(GRK5)–Arr2 from the pACKR3(GRK5)–Arr2–Fab7 complex (PDB entry XXXX) with Fab7 density omitted. (B) Sequence alignment of the arrestin finger and back loops. Conserved hydrophobic residues involved in receptor and detergent/membrane binding are highlighted in green. (C) A flag pulldown assay in the absence or presence of increasing concentrations of CID24 shows no competition with Arr2 binding. CID24 binds to the cytoplasmic cleft of ACKR3 (PDB entry 7SK6) and thus blocks the access to the TM core. (D) Interactions of the pACKR3(GRK5) C-tail with the Arr2 N-lobe in the pACKR3(GRK5)–Arr2–Fab7 complex (PDB entry XXXX). Electron density for the pACKR3(GRK5) phospho-peptide is shown as a wire cage contoured at 12σ. Phosphate interactions below 4 Å are shown as black dash lines. (E) Comparison of Arr2 from the pACKR3(GRK5) structure with the NTSR1 (PDB entry 6UP7) and the M2R (PDB entry 6U1N) complexes after alignment of the receptor TM cores.

A striking feature of the pACKR3(GRK5)–Arr2 complex is that the Arr2 finger loop is not bound in the receptor core but instead inserts into the micellar boundary near TM1 and TM7 of ACKR3 (**Figure 3A**). This is in stark contrast to all other GPCR−arrestin core complexes (rhodopsin, NTSR_1_, M_2_R, β_1_AR, V_2_R, and 5-HT_2A_ serotonin receptor)(Bous et al., 2022; Cao et al., 2022; Huang et al., 2020; Kang et al., 2015; Lee et al., 2020; Staus et al., 2020; Yin et al., 2019; Zhou et al., 2017). The finger loop contains several highly conserved hydrophobic residues (**Figure 3B**) that likely facilitate this interaction. The side chain of Phe244 in the back loop of Arr2 is also positioned to engage the micellar boundary, although the density is less clear in this region. As anticipated, the C-edge loop of Arr2 (residues 191-196) directly inserts into the micellar boundary (**Figure 3A**) as it does in other reported GPCR−Arr2 complexes (Bous et al., 2022; Cao et al., 2022; Huang et al., 2020; Lee et al., 2020; Staus et al., 2020; Yin et al., 2019), the exception being “tail only” complexes (Nguyen et al., 2019). The only region of Arr2 that interacts with the ACKR3 TM core is β17 residues 245-250 which pack against membrane facing residues of TM1 and 7 and likely ICL1, but the micellar boundary obscures details in this region (**Figure 3A**).

Because the structure was strikingly different from what was expected, we tested whether CID24, a Fab that fully occupies the cytoplasmic cleft of activated ACKR3 (**Figure 3C**) (Yen et al., 2022), can displace Arr2. Our *in-vitro* pulldown assays showed that even a 20-fold molar excess of CID24 does not compete with Arr2 binding to pACKR3 (**Figure 3C**), and a consistent amount of CID24 was pulled down with the pACKR3−Arr2 complex in each condition. In contrast, CID24 efficiently blocks GRK2/5 phosphorylation of ACKR3 (**Figure S7**). Thus, Arr2 does not bind to the unoccupied ACKR3 cytoplasmic cleft, but GRKs are dependent on it.

Therefore, Arr2 seems to sense ACKR3 activation solely by interaction with the phosphorylation barcode. Residues 337-343 of the ACKR3 phosphorylated C-tail bind within the N-lobe groove of Arr2, which contain basic residues well known to interact with phosphopeptides (**Figure 3D**). Although the receptor tail density is relatively poor, implying either dynamic or heterogeneous binding/phosphorylation, a reasonable fit was obtained by orienting phospho-Thr338 such that its phosphate is engaged by Arr2-Lys11 and -Arg25 and Arr2-Lys294 in the gate loop. These interactions further mandate that phospho-Thr341 interacts with Arr2-Lys10 and -Lys107. In addition, the side chains of ACKR3-Leu340 and -Leu343 make hydrophobic contacts with residues in β1 and the αN helix of Arr2, thereby serving as hydrophobic anchors. Thus, although Arr2 engages the C-tail of ACKR3 in a canonical manner, the overall configuration is dramatically different relative compared to previously reported structures (**Figure 3E**).

### GRK2 and GRK5 Phosphorylation Barcodes Lead to Distinct Arrestin Binding Behavior

To examine whether different phosphorylation patterns on the C-tail of ACKR3 trigger distinct Arr2 binding modes, we compared the reconstructions of pACKR3(GRK5)–Arr2 and pACKR3(GRK2)–Arr2. The TM helixes of GRK5 phosphorylated ACKR3 are well defined (**Figure 2A, 3A**), whereas those of GRK2 phosphorylated ACKR3 are not (**Figure 2A**). Moreover, 2D class averages of pACKR3(GRK2)–Arr2 indicate heterogeneous binding modes (**Figure 4A**), suggesting that the interface between pACKR3(GRK2) and Arr2 is more dynamic than that of pACKR3(GRK5). To confirm this observation in solution, we labelled the Arr2-V70C and Arr2-L338C variants with monobromobimane (mBrB), which installs the fluorophore into the finger and C-edge loops, respectively, and then assessed whether there are differences when the finger loop or the C-edge loop interacts with pACKR3(GRK2) and pACKR3(GRK5) (**Figure 4B, C**). Both complexes led to an increase in the fluorescence of Arr2-V70CmBrB and Arr2-L338CmBrB. However, pACKR3(GRK5) induced greater fluorescence changes in Arr2-V70mBrB and Arr2-L338CmBrB than pACKR3(GRK2), suggesting that both the Arr2 finger loop and the C-edge loop in pACKR3(GRK5)–Arr2 interact with micelles more efficiently than Arr2 in pACKR3(GRK2)–Arr2. These data are consistent with the observed variability in the cryo-EM class averages with GRK2 phosphorylation (**Figure 4A**).

**Figure 4.**
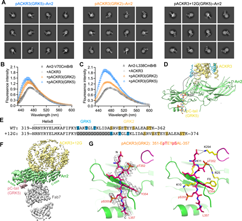
GRK barcoding dictates distinct arrestin binding modes to ACKR3. (A) 2D class averages for pACKR3(GRK5)–Arr2–Fab7, pACKR3(GRK2)–Arr2–Fab7 and pACKR3+12G(GRK5)–Arr2–Fab7. (B, C) Fluorescence spectra of Arr2-V70CmBrB (B) and Arr2-L338CmBrB (C) alone (*black*), or in the presence of non-phosphorylated (*grey*), GRK2 (*orange*) or GRK5 (*blue*) phosphorylated ACKR3. Error bars represent S.E. from three technical replicates. (D) An 18-residue disordered linker from ACKR3 TM7 to the beginning of the GRK5 barcode is shown as a dashed line. The model shown is that of the pACKR3(GRK5)–Arr2–Fab7 complex (PDB entry XXXX). (E) Twelve glycine residues were inserted between Helix8 and the C-tail of ACKR3 (ACKR3+12G) to extend the GRK5 phosphorylation sites to where the GRK2 phosphorylation sites begin. (F) Sharpened map of the 3.3Å pACKR3+12G(GRK5)–Arr2–Fab7 complex highlights a more heterogenous interface between ACKR3+12G and Arr2. (G) Interactions of the ACKR3 C-tail phosphorylated by GRK2 with Arr2 N-lobe in the pACKR3(GRK2)– Arr2–Fab7 complex (PDB entry XXXX). The electron density of ACKR3 GRK2 phospho-peptide is shown as a wire cage contoured at 10σ. Phosphate contacts below 4 Å are shown as black dashed lines.

Besides the distinct barcoding provided by the GRKs (discussed below), the distance between the barcodes and the TM core of ACKR3 is the largest difference between the two structures. There are 18 disordered amino acids between the last modeled residue of TM7 (Ile318) and the installed GRK5 barcode (residues 337-343). The path of the intervening peptide (∼70 Å based on Cα-Cα distances of 3.8 Å, but much less if Helix8 is still intact) would have to circumvent the finger and gate loops and thus follow a path similar to the vasopressin receptor phosphopeptide bound to Arr2 (PDB entry 4JQI) (Shukla et al., 2013) (**Figure 4D**). This suggests that Arr2 would be constrained much closer to the receptor and micelle surface when binding to the GRK5 barcode than the GRK2 barcode, which begins 12 residues more C-terminal (**Figure 4E**). With this additional spacer, Arr2 could readily adopt multiple interaction modes, either binding just to the phosphorylated C-tail (tail only) or making additional micelle contacts with its various membrane binding elements.

To test this idea experimentally, we inserted a flexible linker containing twelve glycine residues between Helix8 and the beginning of the C-tail of ACKR3 (ACKR3+12G) to extend the GRK5 barcode to where the GRK2 barcode begins (**Figure 4E**). This variant is efficiently phosphorylated by GRK5 and forms a stable complex with Arr2, but the 2D class averages indicate heterogeneous interactions with the receptor and there is no well-defined interface between ACKR3 and Arr2 in the best 3D reconstruction (**Figure 4A, F, S8**). Thus, the distance of the barcode with respect to the TM core of the receptor can result in very different configurations of Arr2.

Although the primary sequence and phosphorylation sites are different, the pACKR3(GRK2) C-tail binds to Arr2 in a similar conformation as that of pACKR3(GRK5) (**Figure 3D, 4G**). Phospho-Thr351 interacts with Arr2-Lys11 and - Arg25, phospho-Ser354 interacts with Arr2-Lys10, and the side chains of ACKR3-Tyr353 and -Leu357 form hydrophobic anchors with β1 and αN of Arr2. The same interactions for the ACKR3 C-tail were observed in the nanodisc structure, which were among the most well defined among all our structures (**Figure S9A**). Thus, Arr2 binding to ACKR3 seems to favor a consensus sequence of pTXý(pS/pT)Xý (where ý is hydrophobic and X is any amino acid) although it is not known if both phosphosites in the hexamer are fully modified in our complexes.

### Arr3 and Arr2 bind to ACKR3 differently, but their response to different barcodes is similar

To understand if there are significant differences in how Arr2 and Arr3 bind to GRK-phosphorylated ACKR3, we also determined reconstructions of pACKR3(GRK5)–Arr3 and pACKR3(GRK2)–Arr3 (**Figure 2A, S3, 4**). In the structure of pACKR3(GRK5)–Arr3, the receptor helices could not be clearly resolved in the micelle, and the C-lobe of Arr3 is less ordered (**Figure 2A, 5A**). This is not surprising because Arr3 does not contain one of the membrane-anchoring C-edge loops in its C-lobe (**Figure 5D**). However, the finger loop of Arr3 still inserts into the micelles like that of Arr2 (**Figure 5A**), consistent with the isoforms having a similar set of hydrophobic residues (**Figure 3B**). The lack of the C-edge anchor results in a ∼30° difference in the orientation of Arr2 and Arr3 relative to the detergent micelle surface (**Figure 5B, C**). This suggests that Arr2 and Arr3 could engage ACKR3 in distinct orientations in cells. Because the C-lobe of Arr3 is not fixed relative to the detergent/membrane surface, Arr3 complexes are expected to be more intrinsically flexible, which may be important for its proposed scaffolding roles in other receptors (Peterson and Luttrell, 2017).

**Figure 5.**
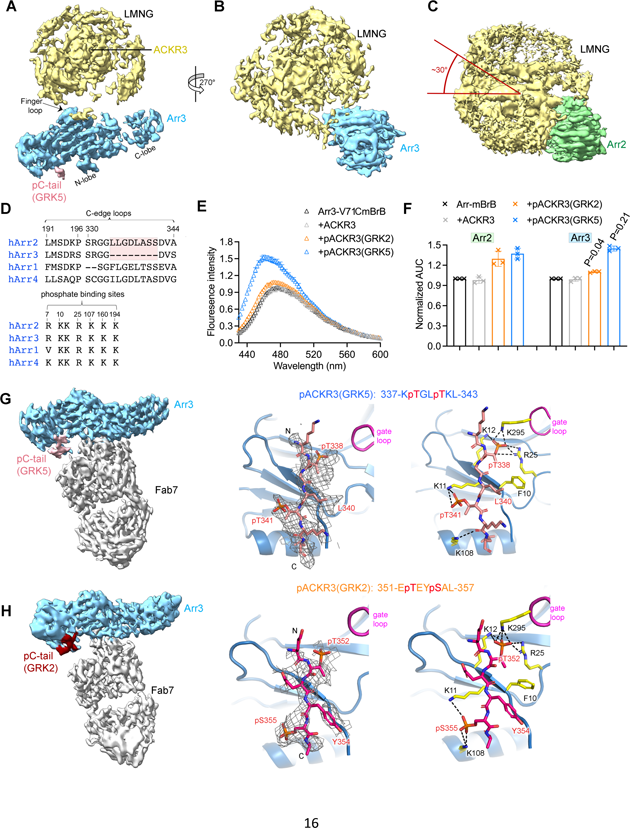
Arr3 binds to ACKR3 in a unique way compared to Arr2, but with similar responses to barcoding by different GRK isoforms. (A, B) Sharpened map of pACKR3(GRK5)–Arr3 from the pACKR3(GRK5)–Arr3–Fab7 complex with Fab7 omitted. (C) Sharpened map of pACKR3(GRK5)–Arr2 from the pACKR3(GRK5)–Arr2–Fab7 complex with Fab7 omitted. Arr2 is positioned at the same angle as Arr3 in (B) to highlight the ∼30° difference in orientation with respect to detergent micelle. (D) Sequence alignment of arrestin C-edge loops and receptor phosphates binding sites. The C-edge loop which is unique in Arr2 is highlighted in pink. (E) Fluorescence spectra of Arr3-V71CmBrB alone (black), or in the presence of non-phosphorylated ACKR3 (grey), pACKR3(GRK2) (orange) or pACKR3(GRK5) (blue). Error bars represent S.D. from three technical replicates. (F) Area under curve (AUC) value from the data in 4B and 5C normalized to Arr2 and Arr3, respectively, allows a direct comparison between Arr2 and Arr3. The AUC value obtained in the presence of GRK2 or GRK5 phosphorylated ACKR3 was compared between Arr2 and Arr3 using t test and p value is shown. (G) Sharpened map of pACKR3(GRK5)–Arr3– Fab7. Interactions of the ACKR3 C-tail phosphorylated by GRK5 with the Arr3 N-lobe in the pACKR3(GRK5)–Arr3– Fab7 complex. Electron density of the pACKR3(GRK5) phospho-peptide is shown as a wire cage contoured at 10σ. Distances below 4Å are shown as black dash line. (H) Sharpened map of pACKR3(GRK2)–Arr3–Fab7. Interactions of the ACKR3 C-tail phosphorylated by GRK5 with the Arr3 N-lobe in the pACKR3(GRK2)–Arr3–Fab7 complex. Electron density of the pACKR3(GRK2) phospho-peptide is shown as a wire cage contoured at 12σ. Distances below 4 Å are shown as black dash line.

In the pACKR3(GRK2)–Arr3 complex, the interface between ACKR3 and Arr3 is the most dynamic and yielded the 3D reconstruction with the lowest resolution (∼7 Å) (**Figure 2A, S4**). In addition to the lack of the C-edge membrane anchor, the longer linker between the GRK2 phosphorylation sites and the receptor core apparently results in a loose interface that primarily features the tail interaction (**Figure 2A, S5**). To further compare the role of finger loops in Arr2 and Arr3 as a function of barcoding, we labelled the Arr3-V71C variant, which installs mBrB into the finger loop. The changes in fluorescence follow similar patterns as Arr2-V70CmBrB (**Figure 5E** versus **Figure 4B**) in that both GRK2 and GRK5 phosphorylated ACKR3, but not non-phosphorylated ACKR3, showed an increase in fluorescence, and pACKR3(GRK5) exhibited a greater enhancement. Interestingly, pACKR3(GRK2) induced much less change in fluorescence in Arr3-V71CmBrB than Arr2-V70CmBrB (**Figure 5F)**, consistent with less extensive interaction between Arr3 and pACKR3(GRK2) embedded micelles, as expected from our cryo-EM analysis (**Figure 2A**). In contrast, pACKR3(GRK5) induced comparable changes in Arr3-V71CmBrB as Arr2-V70CmBrB (**Figure 5F**), which suggests that the finger loop in Arr2 and Arr3 serves a similar role in membrane anchoring.

Despite higher heterogeneity, we could isolate a class of particles centered on Arr3 and Fab7 for close examination of the C-tail interactions with Arr3 (**Figure 5G, H, S3, 4**). Once again, the GRK2 and GRK5 phosphorylated C-tails adopt similar configurations when bound to Arr3 and engage the same residues (**Figure 5G, H**). Because the phosphate binding residues are highly conserved between Arr2 and Arr3 (**Figure 5D**), it is not surprising that the tail interactions are comparable between the two arrestin isoforms, with the same hexameric sequences bound for each GRK barcode.

### Conformational Landscape of Arrestins

As scaffolding proteins, arrestins can achieve selective binding with downstream effectors and thereby modulate cellular responses in several ways, including conformational changes of their effector binding sites, changes in binding site alignment by interdomain twist, and adjustment of accessibility of the sites by adopting different configurations at the receptor and membrane surface (Chen et al., 2018). Although the differences in overall configuration of the pACKR3–Arr2/3 complexes due to the distinct GRK barcoding were profound, differences in arrestin conformation as a function of barcode were more subtle: the RMSD of Arr2 in the pACKR3(GRK2) vs pACKR3(GRK5) complexes was 0.67 Å (based on 302 Cα positions) and for Arr3 it was 0.45 Å (based on 282 Cα positions) (**Figure S9B, C**).

However, comparisons of arrestin conformations in general is potentially complicated from bias introduced by stabilizing Fabs (Fab7 in this study, Fab30 in others) or crystal lattice contacts. To better understand how our structures map in the conformational space of all previously deposited arrestin structures and the potential effects of the Fabs, we used principal component analysis (PCA) (**Figure 6A, S10**). The two largest conformational variances were PC1 and PC2. PC1 (∼79% of the conformational variance among deposited arrestin structures) corresponds to the well-known twist observed upon arrestin activation, such as by disruption of its polar core (**Movie S1**). PC2 (∼8.9% of the variance) corresponds to more of a “wag” of the C-lobe relative to the N-lobe (**Movie S2**). Structures with PC1 values over 15 generally correspond to activated arrestins.

**Figure 6.**
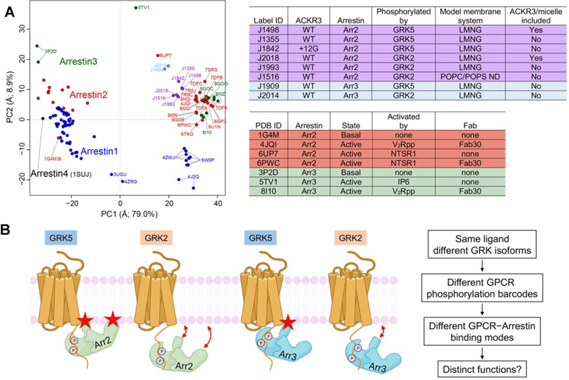
PCA reveals the conformational landscape of arrestins and their complexes with ACKR3. (A) Conformational map derived from all previously deposited arrestin structures with new structures of arrestin–Fab7 complexes from this paper (light blue and purple circles) superposed. Blue, red, green, and black circles otherwise correspond to structures that include Arr1, Arr2, Arr3, and Arr4, respectively. Structures with Fab7 (light blue and purple circles) or Fab30 (red and green and PC1>15, on right) fall in distinct clusters. Detailed information on the models used for PCA is provided in **Table S2**. The PC1 axis corresponds to the well-established twist between the N- and C-lobes of arrestin characteristic of activation (**Movie S1**), whereas the PC2 axis corresponds to an activation-independent “wag” of the C-lobe relative to the N-lobe **(Movie S2)**. The “ACKR3/micelle included” column refers to whether the solubilized receptor was included in the reconstruction (*i.e.,* nanodisc (ND) or LMNG micelle). (A) (B) The distinct configurations of ACKR3–arrestin complexes mediated by different GRK barcodes and different arrestin isoforms identified in this paper may be generally applicable to other 7TM receptors and trigger distinct cellular outcomes. Stars indicate the position of the finger and C-edge loops (Arr2 only) as they engage the membrane. The GRK2 barcode in the C tail of ACKR3 is further from the receptor core than that of GRK5, yielding in our experiments a larger proportion of “tail-mode” complexes. In the case of ACKR3, its 100% bias towards arrestin seems to be entirely driven by GRK phosphorylation and not receptor interactions with arrestin.

Receptor complexes with Fab30, which often involve GPCRs that are fused to the vasopressin receptor C-terminus, cluster tightly around PC1 values of 31-40. This could either indicate that the interaction of Fab30 with both the hinge and bound phosphopeptide puts a constraint on the twist between lobes, or simply that many of these structures involve the same vasopressin phosphopeptide interaction with Arr2 (Chen and Tesmer, 2022). Notably, in the NTSR_1_–Arr2 complex without Fab30 (PDB entry 6UP7) (Huang et al., 2020), the PC1 and 2 values are remarkably different from those of the structure of NTSR_1_–Arr2 complex with Fab30 (PDB entry 6PWC) (Yin et al., 2019). This suggests there could be Fab30-mediated bias, although it could also be a consequence of the different C-tails in the complex (the native sequence in the - Fab30 complex vs a vasopressin C-tail fusion in the +Fab30 complex) (Huang et al., 2020; Yin et al., 2019). Our structures with Fab7 show a similarly tight distribution (PC1 values 18-26) with a degree of twist more consistent with that of PDB entry 6UP7 (PC1 value 16), determined without any Fab (Huang et al., 2020). However, further structure determinations will be needed to evaluate if Fab7 is constraining Arr2/3 conformations.

Finally, the PCA analysis indicates that regardless of GRK barcode, the conformations of Arr3 bound to ACKR3 are distinct from the analogous Arr2 complexes (**Figure 6A**). Thus, although a modest difference, such conformational differences could potentially be interpreted by downstream effectors that rely on binding to both lobes of arrestin.

## Discussion

Prior to this study, we anticipated that active ACKR3 would facilitate GRK and arrestin binding in the “canonical” manner by engaging the cytoplasmic pocket, but that heterotrimeric G proteins would be excluded. However here we show that Arr2 and Arr3 instead bind in a noncanonical manner that does not utilize the open cytoplasmic pocket of the receptor. Instead, the finger and back loops engage the micelles (**Figure 3**). There is some limited contact of Arr2 with the exterior of the receptor when phosphorylated by GRK5, but otherwise it is clear that the major driver for arrestin binding to activated ACKR3 is simply the barcode itself.

But what determines the bias? Because chimeric swaps of the ICLs of ACKR3 with loops from the G protein-coupled chemokine receptor, CXCR2, were ineffective in restoring G protein coupling to ACKR3 and eliminating its arrestin bias (Yen et al., 2022), we speculate that the dynamics of active ACKR3 rather than any specific sequence or configuration of its ICLs may be a determining factor. More specifically, it may be that the dynamic nature of the cytoplasmic domain, as observed in our prior ACKR3 structures (Yen et al., 2022), precludes productive contacts between the receptor cytoplasmic cleft and the arrestin finger loop. Nevertheless, ACKR3 still recruits arrestins because of its phosphorylated C-tail and, depending on the barcode location, insertion of the finger loop into the micelle/bilayer as well. Dynamics may also preclude productive contacts with helix5 of G proteins, leading to a lack of G protein coupling for ACKR3. Indeed, previous studies have suggested that not only the nature but also the duration of intermolecular contacts between the receptor pocket and helix5 contribute to coupling specificity (Sandhu et al., 2022). Therefore, the lack of persistent contacts could prohibit G protein coupling in the case of ACKR3. Thus, dynamic exclusion of both arrestin and G proteins from the cytoplasmic pocket but the ability of arrestin to engage the phosphorylated tail and the membrane/micelle may provide the recipe for a 100% arrestin biased receptor. This further implies that arrestin bias is dictated by the interaction of ACKR3 with GRKs. Since GRKs depend on access to the cytoplasmic cleft (**Figure S7**) (Chen et al., 2021), it remains to be determined how GRKs engage the cleft. Finally, it seems likely that the binding mode(s) of arrestins to ACKR3 observed in this study will also occur in other 7TM receptors. By avoiding extensive contacts within the cytoplasmic cleft, it would be much simpler for arrestins to bind to a larger variety of activated 7TM receptors. Indeed, tail mode Arr2 and Arr3 interactions have now also been reported for the M2 muscarinic (ICL3 phosphorylated) and ACKR2 receptors (Maharana et al., 2023a), suggesting that tail mode interactions could be very common across the 7TM superfamily.

This study also unexpectedly revealed that the finger loop of arrestin, which is allosterically altered during arrestin activation, plays a direct role in micelle and potentially membrane binding. As such, our structures provide a molecular mechanism for the recent observation that arrestins can anchor to the plasma membrane via the C-edge and finger loops in the absence of receptor, and that receptor-activated arrestins can remain membrane bound and migrate on the membrane surface even after they dissociate from activated receptors (Grimes et al., 2023). We speculate that in the case of Arr2, the interactions of the membrane with both the finger loop and the C-edge loop (and perhaps the binding of negatively charged phospholipids to the C-lobe) are sufficient to hold arrestins in an active-like conformation well after their polar cores are disrupted by phosphorylated receptor loops or tails.

Differential barcoding by GRKs (or other second messenger kinases) has long been evoked to explain why different GRKs can lead to different cellular outcomes. For example, both GRK2 and GRK6 phosphorylation of the β_2_ adrenergic receptor initiates the desensitization of G protein signaling, but only GRK6 phosphorylation is required for ERK1/2 activation (Nobles et al., 2011). However, until now there has not been a molecular explanation. Here we show that phosphorylation barcoding of ACKR3 by GRK2 and GRK5 leads to dramatically different behavior and dynamics in individual complexes regardless of the bound arrestin isoform, with GRK2 phosphorylation leading to more “tail mode”-like interactions and GRK5 to more compact and rigid assemblies with the arrestin bound to the micelle (**Figure 6B**).

These unique configurations could lead to differences in the persistence of the bound arrestin and/or contribute to distinct downstream outcomes. In fact, in recent studies we showed that GRK2 and GRK5 phosphorylated ACKR3–Arr complexes have substantially different half-lives (Schafer et al., 2023). We also showed a unique twist on the functional role of the GRKs where GRK2 phosphorylation of ACKR3 is operative only when CXCR4 is also present and activated by CXCL12 because the kinase depends on G_βγ_. In contrast, in the absence of CXCR4, GRK5 dominates ACKR3 phosphorylation due to its independence from heterotrimeric G protein activation (Schafer et al., 2023). The impact of the different barcodes is yet to be determined but GRK2 phosphorylation may not only regulate ACKR3 function but also have indirect effects on CXCR4.

We also demonstrated that Arr2 and Arr3 bind to detergent solubilized ACKR3 in markedly different orientations that seem to be dictated by differences in their primary sequence (**Figure 5**). Their ∼30° difference in orientation could also underly distinct downstream events selectively mediated by Arr2 or Arr3. In contrast, we found only small differences in the conformation of Arr2/3 when bound to different GRK barcodes (**Figure 6A**). As noted above, it remains possible that barcode dependent differences in Arr2 or Arr3 in response to different barcodes are somewhat muted by Fab7 binding in the hinge.

Surprisingly, the conformation and interactions of Arr3 in its complexes with pACKR3(GRK2) and pACKR3(GRK5) differ markedly from those of Arr3 in complex with ACKR3 phosphopeptides reported elsewhere (Min et al., 2020; Sarma et al., 2022). Our structures also reveal the expected canonical activated conformation for Arr3 (**Figure 6A**), whereas the Arr3– peptide complexes had intermediate and heterogeneous degrees of domain rotation. Although the difference could be due to the fact that we used full length ACKR3, which could impose structural constraints that phosphopeptides do not, the many structural implausibilities found in the deposited 6K3F structure (Min et al., 2020) prevent us from further speculation. For this reason, we left 6K3F structure out of our PCA analysis (**Figure 6A**), but their mapped PC1 and 2 values are included in the supplementary information for reference (**Table S2**).

In summary, we developed a new tool (Fab7) that stabilizes the active state of Arr2 and 3 and enabled us to explore the molecular consequences of differential GRK barcoding of a receptor in its native form regardless of barcoding (**Figure 6B**). Fab7 will be useful for interrogating arrestin binding to many other receptors in a manner that is likewise not dependent on the barcode. It could also be engineered in the future as an intrabody for pinpointing the location of arrestin activation in living cells or selectively blocking distinct scaffolding activities of Arr2/3. We have also shown that arrestins bind to ACKR3 in a manner that is distinct from other GPCR–Arr complexes, providing insights into the molecular basis for the complete bias of ACKR3 for arrestins. Finally, we have shown that the distinct barcodes lead to different ACKR3–arrestin configurations for a given arrestin, and that Arr2 and Arr3 bind to ACKR3 in fundamentally different ways irrespective of barcode. All of these molecular differences, individually or in sum, may contribute to GRK/arrestin isoform dependent cellular outcomes.

**Figure S1.**
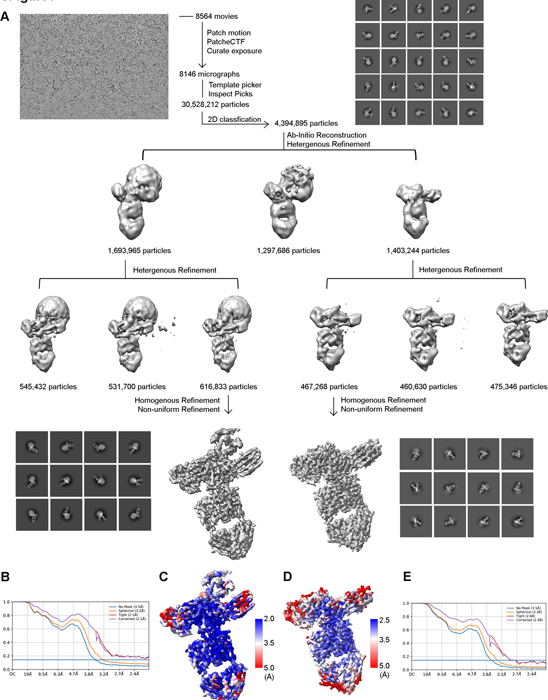
Workflow of cryo-EM data processing and resolution analysis of pACKR3(GRK5)–Arr2–Fab7. (A) Representative micrograph shows well-distributed complexes. The cryo-EM workflow from motion correction to CTF estimation to particle picking to 2D classification to 3D refinement is shown. (B, E) Fourier shell correlation (FSC) curves calculated by cryoSPARC with 0.143 as a cutoff for pACKR3(GRK5)–Arr2–Fab7 with (B) or without (E) the ACKR3 TM core. (C, D) Local resolution estimation calculated by cryoSPARC for pACKR3(GRK5)–Arr2–Fab7 with (A) (C) or without (D) the ACKR3 TM core.

**Figure S2.**
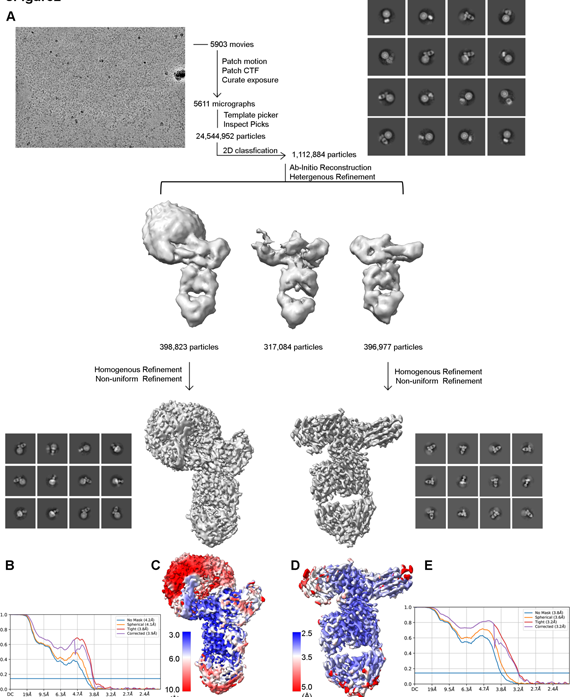
Workflow of cryo-EM data processing and resolution analysis of pACKR3 (GRK2)–Arr2–Fab7. (A) Representative micrograph shows well-distributed complexes. The cryo-EM data processing workflow from motion correction to CTF estimation to particle picking to 2D classification to 3D refinement is shown. (B, E) FSC curves calculated by cryoSPARC with 0.143 as a cutoff for pACKR3(GRK2)–Arr2–Fab7 with (B) or without (E) the ACKR3 TM core. (C, D) Local resolution estimation calculated by cryoSPARC for pACKR3(GRK2)–Arr2–Fab7 with (B) or without (D) the ACKR3 TM core.

**Figure S3.**
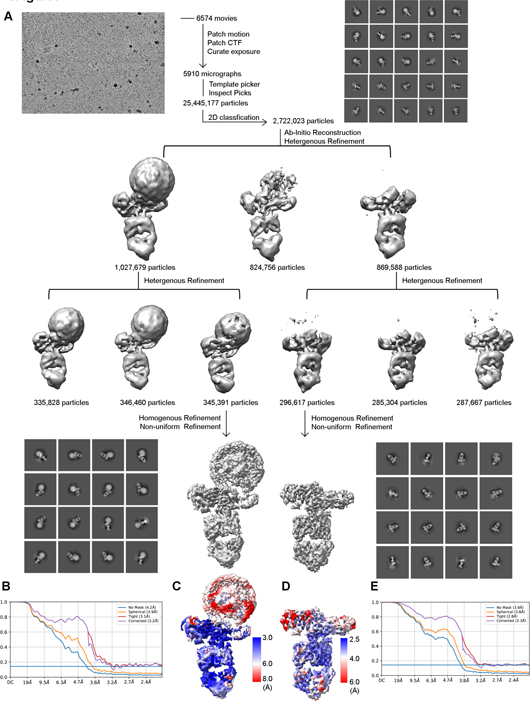
Workflow of cryo-EM data processing and resolution analysis of pACKR3(GRK5)–Arr3–Fab7. (A) Representative micrograph shows well-distributed complexes. The cryo-EM data processing workflow from motion correction to CTF estimation to particle picking to 2D classification to 3D refinement is shown. (B, E) FSC curves calculated by cryoSPARC with 0.143 as a cutoff for pACKR3(GRK5)–Arr3–Fab7 with (B) or without (E) the ACKR3 TM core. (C, D) Local resolution estimation calculated by cryoSPARC for pACKR3(GRK5)–Arr3–Fab7 with (C) or without (D) the ACKR3 TM core.

**Figure S4.**
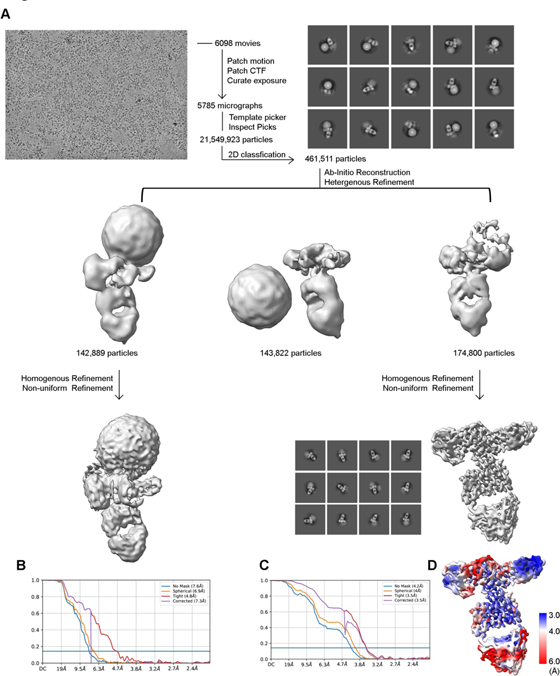
Workflow of cryo-EM data processing and resolution analysis of pACKR3 (GRK2)–Arr3–Fab7. (A) Representative micrograph shows well-distributed complexes. The cryo-EM data processing workflow from motion correction to CTF estimation to particle picking to 2D classification to 3D refinement is shown. (B, C) FSC curves calculated by cryoSPARC with 0.143 as a cutoff for pACKR3(GRK2)–Arr3–Fab7 with (B) or without (C) the ACKR3 TM core. (E) Local resolution estimation calculated by cryoSPARC for pACKR3(GRK2)–Arr2–Fab7 without the ACKR3 TM core.

**Figure S5.**
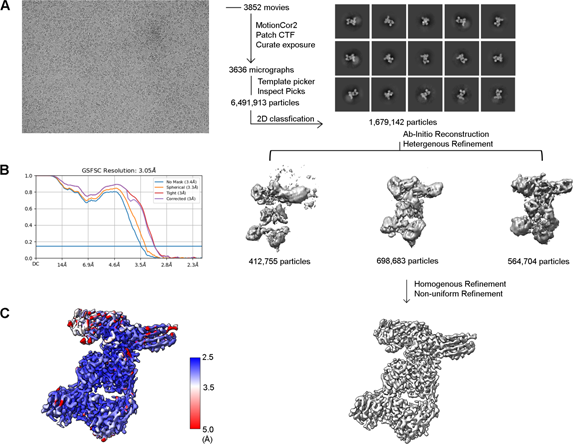
Workflow of cryo-EM data processing and resolution analysis of pACKR3 (GRK2)–Arr2–Fab7 in nanodisc. (A) Representative micrograph shows well-distributed complexes. The cryo-EM data processing workflow from motion correction to CTF estimation to particle picking to 2D classification to 3D refinement is shown. (B) FSC curves calculated by cryoSPARC with 0.143 as a cutoff for pACKR3(GRK2)–Arr2–Fab7 in nanodisc without the ACKR3 TM core. (E) Local resolution estimation calculated by cryoSPARC for pACKR3(GRK2)–Arr2–Fab7 without the ACKR3 TM core.

**Figure S6.**
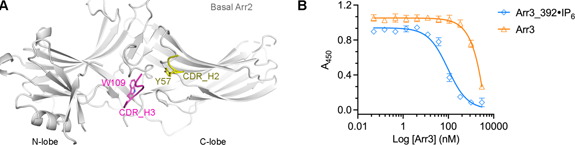
Fab7 preferentially binds arrestin in its activated state. (A) Superposition of basal Arr2 (light grey, PDB entry 1G4M) with activated Arr2 from the pACKR3(GRK2)–Arr2–Fab7 complex (only Fab7 shown, PDB entry XXXX) aligned on their Arr2 C-lobes. Some of the potential clashes expected to reduce affinity are highlighted. (B) ELISA analysis of Fab7 competition assay reveals that preactivated Arr3_392·IP6 (IC50 ∼90 nM) competes for Fab7 binding more efficiently than WT Arr3 (IC50 ∼ 35 µM). Error bars represent S.D. from three technical replicates.

**Figure S7.**
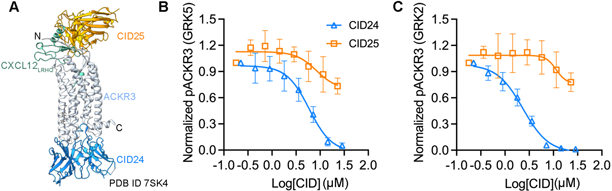
CID24 efficiently blocks GRK2 and GRK5 phosphorylation of ACKR3. (A) CID24 and CID25 bind to the intercellular and extracellular regions of ACKR3, respectively (PDB entry 7SK4). (B, C) ACKR3 phosphorylation by GRK5 (B) and GRK2 (C) in the presence of increasing amounts of CID24 or CID25.

**Figure S8.**
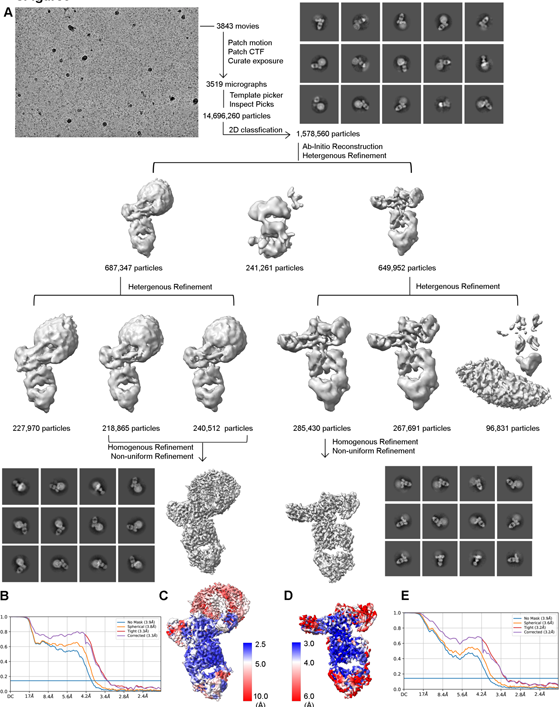
Workflow of cryo-EM data processing and resolution analysis of pACKR3+12G(GRK5)–Arr2– Fab7. (A) Representative micrograph shows well-distributed complexes. The cryo-EM data processing workflow from motion correction to CTF estimation to particle picking to 2D classification to 3D refinement is shown. (B, C) FSC curves calculated by cryoSPARC with 0.143 as a cutoff for pACKR3+12G(GRK5)–Arr2–Fab7 with (B) or without (C) the ACKR3 TM core. (E) Local resolution estimation calculated by cryoSPARC for pACKR3+12G(GRK5)–Arr2–Fab7 without the ACKR3 TM core.

**Figure S9.**
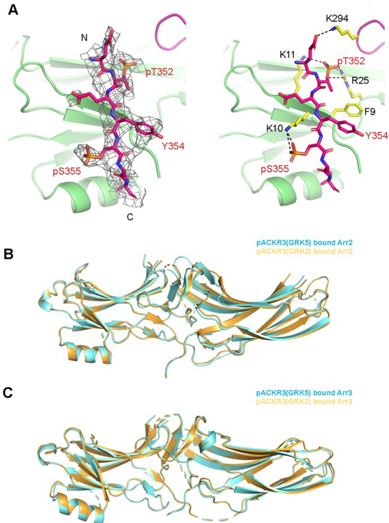
(A) Interactions of ACKR3 C-tail phosphorylated by GRK2 with Arr2 N-lobe in the pACKR3(GRK2)– Arr2–Fab7 complex in nanodisc. Electron density of ACKR3 GRK2 phospho-peptide is shown as a wire cage contoured at 4σ. Distances below 4 Å are shown as black dash line. (B, C) Alignment of pACKR3(GRK5) bound and pACKR3(GRK2) bound Arr2 (B) or Arr3 (C) on the N-lobe suggests subtle changes in arrestin conformation.

**Figure S10.**
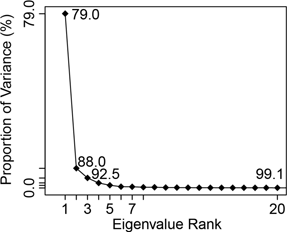
Scree plot showing associated eigenvalues from PCA analysis of deposited arrestin structures (Figure 6A). The eigenvalues measure the conformational variance along corresponding principal component axes, which usually decrease rapidly after the top few components, as occurs here.

**Table S1.**
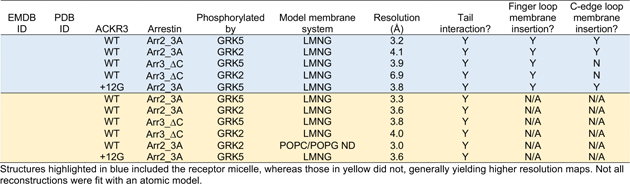
Summary of determined cryo-EM maps and structures.

**Table S2. List of PDB entries used for PCA analysis and their PC1 and PC2 values.**

**Movies S1 & S2**. **Motions described by the PC1 and PC2 axes from PCA analysis, respectively.** The portion of arrestin structure used for PCA is shown as tube with its N- and C-lobes colored green and blue, respectively. Dashed lines indicate loops not used for PCA because of structural gaps among the compared structures. The range of motion is determined by 1.5-fold standard deviation of conformations along the PC in both directions from the mean conformation (PC values of 0). PC1 motion corresponds to a ∼30° twist of the C-lobe relative to the N-lobe from lower to higher PC1 values. PC2 motion corresponds a ∼5° “wag” of the C-lobe relative to the N-lobe.

## CONTACT FOR REAGENT AND RESOURCE SHARING

Please contact Qiuyan Chen, John J. G. Tesmer, or Tracy Handel with any requests regarding reagents used in this study.

## METHOD DETAILS

### Expression and purification of ACKR3

ACKR3 was co-expressed with CXCL12_LRHQ_ in *Sf9* cells as previously described (Yen et al., 2022). Briefly, *Sf9* cells were infected (multiplicity of infection of 6 for each virus) with separate baculoviruses (prepared using the Bac-to-Bac Baculovirus Expression System, Invitrogen) containing the CXCL12_LRHQ_ gene as well as the ACKR3 gene (residues 2-362) with an N-terminal HA signal sequence, and tandem C-terminal 10xHis and FLAG purification tags. After 48 hours, the infected cells were harvested by centrifugation and the membranes prepared by four rounds of dounce homogenization, first in hypotonic buffer containing 10 mM HEPES (pH 7.5), 10 mM MgCl_2_, and 20 mM KCl, followed by three more washes with hypotonic buffer plus 1 M NaCl. The membranes were spun down at 50,000 x g for 30 min and resuspended between each round of douncing. The samples were then solubilized in 50 mM HEPES pH 7.5, 400 mM NaCl, 0.75/0.15% lauryl maltose neopentyl glycol/cholesteryl hemisuccinate (LMNG/CHS) with a protease inhibitor tablet (Roche) for 4 hrs. Insoluble material was removed by centrifugation at 50,000 x g for 30 min. Talon resin (Clontech) with 20 mM imidazole was added to the soluble fraction and incubated overnight at 4 °C. The resin was then transferred to a plastic purification column and washed with washing buffer 1 containing 50 mM HEPES (pH 7.5), 400 mM NaCl, 10% glycerol, and 20 mM imidazole, plus 0.1/0.02% LMNG/CHS, followed by washing buffer 1 plus 0.025/0.005% LMNG/CHS and finally eluted with washing buffer 1 plus 0.025/0.005% and 250 mM imidazole. The imidazole was removed with a desalting column (PD MiniTrap G-25, GE Healthcare). The final protein concentration was determined by A_280_ using an extinction coefficient of 85000 M^-1^cm^-1^, snap frozen in liquid nitrogen, and stored at -80 °C for later use. A similar strategy was used to prepare complexes of ACKR3+12G.

### Expression and purification of GRK5

GRK5 was expressed and purified from *E. Coli* cells as previously described (Beyett et al., 2020). Briefly, a pMAL plasmid containing human full-length GRK5 with a C-terminal 6xHIS tag was transformed into *E. coli* Rosetta cells. The expression of GRK5 was induced with 200 µM IPTG at OD around 0.6-0.8 and the cultures were shaking at 18 °C overnight. For purification, cell pellets were resuspended and homogenized in lysis buffer containing 20 mM HEPES (pH 8.0), 400 mM NaCl, 0.1% Triton-X (v/v), 2 mM DTT, DNase, 0.1 mM PMSF, leupeptin, and lima bean trypsin protease inhibitor. The cells were then lysed using an Avestin C3 emulsifier and centrifuged at 18,000 rpm for 30 min. The supernatant was combined and loaded onto a 3 ml home-packed Ni^2+^-NTA column pre-equilibrated with buffer A containing 20 mM HEPES (pH 8.0), 400 mM NaCl and 0.5 mM DTT. The column was then washed with 50 ml buffer A, followed by 100 ml buffer B containing 20 mM HEPES (pH 8.0), 100 mM NaCl and 0.5 mM DTT plus 20 mM imidazole. The bound protein was eluted with buffer B plus 200 mM imidazole and then loaded onto a linked 1 ml HiTrap Q HP (that it flows through) and 1 ml HiTrap SP HP column (that it binds). The columns were then uncoupled and a linear NaCl gradient (0.1-0.6 M) was used to elute GRK5 from the SP column. GRK5 elutes with ∼0.3-0.5 M NaCl. The fractions containing GRK5 were combined, concentrated with a 50 kDa cutoff Amicon concentrator to ∼500 µl, then further purified using a Superdex 200 Increase 10/300 GL column equilibrated with 20 mM HEPES (pH 8.0), 100 mM NaCl, and 0.5 mM TCEP. The peak fractions were collected, concentrated with a 50 kDa cutoff Amicon concentrator, and stored at -80°C.

### Expression and purification of GRK2

GRK2 S670A was expressed and purified from *Sf9* cells as previously described (Schafer et al., 2023). Briefly, human GRK2 S670A with a C-terminal 6xHIS tag was expressed using the Bac-to-Bac insect cell expression system (Life Technologies). The insect cells were harvested 48 hours post-infection and homogenized with buffer containing 20 mM HEPES (pH 8.0), 400 mM NaCl, 2 mM DTT, 1 mM PMSF, leupeptin, and lima bean trypsin protease inhibitor. The cells were lysed using an Avestin C3 emulsifier and clarified by centrifugation at 35,000xg for 60 min. GRK2 was purified using immobilized metal ion affinity chromatography as described above for GRK5. The purity of GRK2 after this step was ∼90%. Fractions containing GRK2 were pooled and further purified on a Superdex 200 Increase 10/300 GL column equilibrated with 20 mM HEPES (pH 8.0), 100 mM NaCl, and 0.5 mM TCEP. The peak fractions were collected, concentrated with a 50 kDa cutoff Amicon concentrator, and stored at -80°C.

### Expression and purification of Arrestins

Expression and purification of Arr2/3 from *E. Coli* cells was described previously (Vishnivetskiy et al., 2014). WT or variants of Arr2 and Arr3 were prepared using the same procedure. Briefly, the pTrcHisB plasmid containing bovine Arr2 or Arr3 was transformed into *E. coli* Rosetta cells and protein expression was induced with 25 µM (Arr2) or 37.5 µM (Arr3) IPTG for 4 hours at 30 °C. The cell pellets were resuspended and homogenized in buffer containing 20 mM MOPS (pH 7.5), 400 mM NaCl, 5 mM EDTA, 2 mM DTT, 1 mM PMSF, leupeptin, and lima bean trypsin protease inhibitor. Cells were lysed using an Avestin C3 emulsifier and the lysate was clarified by centrifugation at 18,000 rpm for 30 min. The supernatant was collected and arrestin was precipitated by the addition of (NH_4_)_2_SO_4_ to a final concentration 0.32 mg/ml. Precipitated arrestin was collected by centrifugation at 18,000 rpm for 90 min. The pellet was then dissolved in buffer containing 20 mM MOPS (pH 7.5), 2 mM EDTA, and 1 mM DTT, and then centrifuged at 18,000 rpm for 60 min to remove insoluble parts. The supernatant containing soluble arrestin was applied to a heparin column and eluted with a linear NaCl gradient (0.2-1 M). Fractions containing arrestin were identified by SDS-PAGE and combined. For Arr2, the salt concentration of the pooled fractions was adjusted to 50 mM, loaded onto a 5 ml HiTrap Q HP column (Cytiva), and eluted with a linear NaCl gradient. For Arr3, the salt concentration of the pooled fractions was adjusted to 100 mM, and the solution was loaded onto a linked 1 ml HiTrap Q HP (that it flows through) and 1 ml HiTrap SP HP column (that it binds). The columns were then uncoupled and a linear NaCl gradient (0.2-1 M) was used to elute Arr3 from the SP column. The fractions containing arrestin were concentrated with a 30 kDa cutoff Amico concentrator to ∼500 µl, then further purified using a Superdex 200 increase 10/300 GL column (Cytiva) equilibrated with 20 mM MOPS (pH 7.5), 150 mM NaCl, and 0.5 mM TCEP. The peak fractions were collected, concentrated with a 30 kDa cutoff Amicon concentrator, and stored at -80 °C.

### Phosphorylation of ACKR3

Phosphorylation conditions were optimized to achieve maximum phosphorylation of ACKR3. For GRK5, the phosphorylation reaction contained 2 µM ACKR3, 1 µM GRK5, 20 µM c8-PIP_2_ and 200 µM ATP in phosphorylation buffer containing 20 mM HEPES (pH 8.0), 10 mM MgCl_2_ and 0.002% LMNG. Everything except ATP was mixed first and incubated at room temperature for 20 min. ATP was added to initiate phosphorylation and the reaction allowed to proceed for one hour at room temperature. For GRK2, the phosphorylation reaction contained 2 µM ACKR3, 3 µM GRK2 and 200 µM ATP in phosphorylation buffer containing 20 mM HEPES (pH 8.0), 10 mM MgCl_2_ and 0.02/0.002% LMNG/CHS. The phosphorylation reaction was incubated overnight at room temperature.

### Arrestin pulldown

ACKR3 (2 µM) was phosphorylated by GRK5 or GRK2 using the procedure described above. Different arrestin variants (2.4 µM) and Fab7 (2.4 µM) were added to the ACKR3 phosphorylation reaction and incubated for one hour at room temperature. Anti-Flag M2 magnetic beads (Sigma) were washed with buffer A containing 20 mM HEPES (pH 8.0), 100 mM NaCl, 0.01/0.001% LMNG/CHS and then added to the mixture. The sample tubes were incubated in a rotator for an hour at room temperature. Anti-Flag M2 beads were washed five times with 1 ml buffer A and then eluted with buffer A supplemented with 3xFlag peptide (Sigma). Unphosphorylated ACKR3, used as a control, was prepared by omitting ATP in the first phosphorylation step. The eluted samples were analyzed by SDS-PAGE gel electrophoresis with Coomassie staining. The densities of bound arrestin and ACKR3 were quantified using Image Lab and the ratios between different samples were compared.

### Trypsin digestion of arrestin

WT Arr2 (10 µM) was incubated with 2-fold serial dilutions of 32 µM Fab7 at room temperature for 20 min. Trypsin (0.0015 mg/ml) was added to the reaction and the digestion was incubated for 10 min at room temperature. SDS loading buffer was used to quench the reaction. The samples were analyzed by SDS-PAGE gel electrophoresis with Coomassie staining.

### Monobromobimane labeling of arrestin

Arr2-V70C, -L338C and Arr3-V71C with all native cysteines mutated were prepared as described above in buffer containing 20 mM MOPS (pH 8.0) and 150 mM NaCl. A twenty-molar excess of freshly prepared mBrB (Sigma) was added to the reaction and incubated on ice overnight. The sample was loaded on a Superdex 200 Increase 10/300 GL column equilibrated with buffer containing 20 mM MOPS (pH 8.0) and 150 mM NaCl to get rid of excess mBrB. The fractions containing mBrB-labelled arrestins were collected, concentrated with a 30 kDa cutoff Amicon concentrator, and stored at -80 °C. The mBrB labeling level was estimated to ∼75% based on the mBrB fluorescence absorption at 383 nm and the protein absorption at 280 nm.

### Fluorescence measurements

The mBrB-labelled Arr2-V70C, -L338C or Arr3-V71C (2 µM) was incubated with 2 µM ACKR3 and 2 µM ACKR3 phosphorylated by GRK2 or GRK5 at room temperature for 20 min. The reactions were transferred to a 384 black clear bottom plate (Corning) for fluorescence measurement using a plate reader (BioTek). The excitation wavelength was set to 375 nm and the absorption was monitored from 420 nm to 700 nm.

### Preparation of biotinylated Arr3

Arr3_ΔC (1 mg/ml) was incubated with 200 µM IP_6_ for 20 min in buffer containing 20 mM MOPS (pH 8.0), 150 mM NaCl, and 0.5 mM TCEP. A twenty-molar excess of freshly prepared EZ-link sulfo-NHS-Biotin (Thermo fisher) was added to the reaction and incubated on ice for 2 hours. The sample was loaded onto a Superdex 200 Increase 10/300 GL column equilibrated with buffer containing 20 mM MOPS (pH 8.0), 150 mM NaCl, 0.3 mM TCEP and 0.2 mM IP_6_ to remove excess sulfo-NHS-Biotin. Arr3_ΔC biotinylated in the presence of IP_6_ eluted at the retention volume corresponding to a trimer and the peak fractions were collected for Fab selection. The biotinylation level was estimated to be >95% based on pulldown assays using avidin agarose beads.

### Phage display selections

Biotinylated Arr3_ΔC loaded with IP_6_ was used for phage display selection. Phage display selection was performed at 4 °C according to published protocols (Paduch et al., 2013). The selection buffer contained 20 mM MOPS (pH 8.0), 150 mM NaCl, 0.01% LMNG, 0.5% BSA and 0.2 mM IP_6_. In brief, for the first round of selection, 200 nM of biotinylated Arr3_ΔC·IP_6_ complex was immobilized on 250 µl Streptavidin magnetic beads (Promega, Cat No: Z5482) and incubated with 100 μl of a phage library E (Miller et al., 2012) containing 10^12^ phage for 30 min. The resuspended beads containing bound virions were washed extensively and then used to infect freshly grown log phase *E. coli* XL1-Blue cells. Phages were amplified overnight in 2xYT media with 50 µg/ml ampicillin and 10^9^ p.f.u./ml of M13-KO7 helper phage. To increase the stringency of selection, four additional rounds of sorting were performed with decreasing target concentration in each round (second round, 50 nM; third round, 50 nM; fourth and fifth round, 10 nM) using the amplified pool of phage from the preceding round as the input. Selection from the second to fifth rounds was done on an automated Kingfisher automated purification instrument (Thermo Scientific) where the target was premixed with the amplified phage pool and then Streptavidin beads were added to the mixture. From the second round onwards, the bound phages were eluted using 0.1 M glycine (pH 2.7). To eliminate the non-specific and Streptavidin binders, the precipitated phage pool from the second round onwards were negatively selected against 100 µl of Streptavidin beads before adding to the target. The pre-cleared phage was then used as an input for the selection.

### Single-point phage ELISA

All ELISA experiments were performed at 4°C in 96-well plates coated with 50 µl of 2 µg/ml neutravidin in Na_2_CO_3_ buffer (pH 9.6) and subsequently blocked by 1% BSA in PBS. A single-point phage ELISA was used to rapidly screen the binding of the obtained clones. Colonies of *E. coli* XL1-Blue harboring phagemids from 4^th^ and 5^th^ rounds of selection were inoculated directly into 500 μl of 2xYT broth supplemented with 100 μg/ml ampicillin and M13-KO7 helper phage. The cultures were grown overnight at 37 °C in a 96-deep-well block plate. The phage display selection buffer contained 20 mM MOPS (pH 8.0), 150 mM NaCl, 0.01% LMNG, 0.5% BSA and 0.2 mM IP_6_. Culture supernatants containing Fab phage were diluted tenfold in the selection buffer. After 15 min of incubation, the mixtures were transferred to ELISA plates previously incubated with 40 nM biotinylated Arr3_ι1C in experimental wells and with buffer in control wells for 15 min. The ELISA plates were incubated with the phage for another 15 min and then washed with ELISA buffer. The washed ELISA plates were incubated with a 1:1 mixture of mouse anti-M13 monoclonal antibody (cat: 27-9420-01, GE, 1:5,000 dilution in ELISA buffer) and peroxidase conjugated goat anti-mouse IgG (cat: 115-035-003, Jackson Immunoresearch, 1:5000 dilution in ELISA buffer) for 30 min. The plates were washed again, developed with 3,3’,5,5’-Tetramethyl-benzidine/H2O2 peroxidase substrate (TMB) (Thermo Scientific, Cat No: 34021) and then quenched with 1.0 M HCl, and the absorbance at 450 nm was read on a plate reader. Phagemid DNA from the clones from wells with high signal/noise ratio were sequenced to identify the unique binders.

### Sequencing, cloning, overexpression and purification of Fab fragments

The sequencing, cloning, overexpression and purification of the Fab fragments were performed according to published protocols (Bloch et al., 2021).

### Multipoint ELISA for EC50 determination

Multipoint ELISA assays were performed at 4°C to estimate the affinity of the Fabs for Arr3_ι1C. The phage display selection buffer containing 20 mM MOPS (pH 8.0), 150 mM NaCl, 0.01% LMNG, 0.5% BSA and 0.2 mM IP_6_ was used as ELISA buffer. 40 nM of biotinylated target immobilized on a neutravidin coated ELISA plate was incubated with 3-fold serial dilutions of 4 μM purified Fabs for 20 min. The plates were washed, and the bound target-Fab complexes were incubated with an HRP-conjugated Pierce recombinant protein L (cat: 32420, Thermofisher, 1:5000 dilution in ELISA buffer) for 30 min. The plates were washed again, developed with TMB and quenched with 1.0 M HCl, and phage quantified by the absorbance at 450 nm. EC_50_ values were determines by fitting the data with a dose response sigmoidal function in GraphPad PRISM.

### Fab7 competition assays

A multipoint ELISA assay (described above) was used to determine that 50 nM Fab7 reached 50–70% of maximum binding to 25 nM biotinylated Arr3_ι1C •IP_6;_ thus, 50nM was subsequently used in competition assays. For these experiments, 50 nM Fab7 was incubated separately with 3-fold serial dilutions of 3 μM competitors (Arr2, Arr2_3A, Arr3, or Arr3_392) for 30 min. The samples were then transferred to ELISA plates containing 25 nM biotinylated Arr3_ΔC •IP_6_ and incubated for 15 min to capture free Fab7. The plates were then washed with the ELISA buffer, and the bound Arr3_ΔC•IP_6_–Fab7 complexes incubated with HRP-conjugated Pierce recombinant protein L (cat: 32420, Thermofisher, 1:5000 dilution in ELISA buffer) for 30 min. The plates were again washed, developed with TMB and quenched with 1.0 M HCl, and Fabs quantified by A_450_. IC_50_ values were calculated using GraphPad PRISM.

### Cryo-EM sample preparation and image acquisition

Flag pulldown assays described above were used to prepare ACKR3–arrestin–Fab7 complexes for cryo-EM. Quantifoil R1.2/1.3 300-mesh Cu grids were glow-discharged using EasiGlow at 25 mA for 60s. Purified ACKR3–arrestin–Fab7 (3.3 μl at ∼0.6 mg/ml) was applied to the grids and the grids blotted with filter paper for 3.5 s before being plunge-frozen in liquid ethane using a Vitrobot MK IV (Thermo Fisher Scientific). Data were collected on a Titan Krios G4 electron microscope (FEI) equipped with a post-GIF K3 direct electron detector (Gatan) and a Quantum GIF energy filter (Gatan) in the Purdue Life Sciences Cryo-EM Facility. Micrographs were collected in super-resolution mode with a pixel size of 0.527 Å, at a defocus range of 0.6 to 2.5 μm using EPU, and 40 frames were recorded for each movie stack at a frame rate of 78 milliseconds per frame and a total dose of 53.8 electrons/Å^2^.

### Cryo-EM data processing

Cryo-EM movies were imported to cryoSPARC and processed using the standard workflow (Punjani et al., 2017). Beam-induced motion was corrected and binned twofold using Patch Motion in cryoSPARC. The contrast transfer function (CTF) parameters were estimated using the Patch CTF module. Blob picker was used to pick particles on a small set of micrographs to generate class averages as templates for subsequent autopicking using template picker. Several rounds of 2D classification were performed to exclude bad particles that fell into 2D averages with poor features. Particles from different views were selected to generate three initial models using ab initio reconstruction. The resulting 3D models were used for heterogeneous refinement in cryoSPARC. Another round of heterogenous refinement was performed to select 3D classes showing the highest-resolution features for pACKR3(GRK5)–Arr2–Fab7, pACKR3(GRK5)– Arr3–Fab7 and pACKR3_12G(GRK5)–Arr2–Fab7. The selected classes were then refined using homogeneous refinement and nonuniform refinement (Punjani et al., 2020). The image processing flowcharts for each dataset are shown in **Figure S1-5, S8**.

### Model building and refinement

A homology model of Fab7 generated using the SWISS-MODEL server (http://swissmodel.expasy.org), the crystal structure of active Arr2 (PDB entry 4JQI) and the anti-Fab hinge-binding nanobody (PDB entry 6WW2) were docked into the highest resolution cryo-EM map (pACKR3(GRK2)–Arr2–Fab7 nanodisc complex, **Figure S5**) using Phenix (Liebschner et al., 2019). The CDR regions of the Fab7 heavy chain were rebuilt manually in COOT (Emsley et al., 2010). The resulting model was further improved using several rounds of real space refinement in Phenix and manual adjustment in COOT. The same strategy was employed to build and refine the rest of the models except that different initial models were used for different maps: the Fab7 model from the cryo-EM structure described above, the crystal structure of active Arr2 (PDB entry 4JQI), the crystal structure of active Arr3 (PDB entry 5TV1), and the CID24-CXCL12_LRHQ_-ACKR3 complex (PDB entry 7SK6). All figures were prepared using PyMOL and ChimeraX (Pettersen et al., 2021).

### Principal Component Analysis (PCA)

PCA was performed on previous experimental arrestin structures using Bio3D (Grant et al., 2006; Grant et al., 2021; Skjærven et al., 2014). New structures not used for PCA were projected (along with the previous structures) onto the PC1-PC2 plane for structural comparisons. A total of 114 structures of Arr1-4 were collected from the PDB (**Table S2**). Sequences of these structures were aligned using MUSCLE (Edgar, 2004). Prior to PCA, structurally invariant “core” residues were identified through iterated rounds of structural superimposition as previously described (Gerstein and Altman, 1995). These core residues, which were all from the N-lobe, were used as the reference for the superimposition of structures. The new models from this work and six different chain models from (PDB entry 3K6F) (Min et al., 2020) were aligned and superimposed in the same way as the base structures. PCA was calculated for aligned positions where no gap was found for any of the structures. Movies showing morphs for the PC1 and PC2 motions (**Movies S1 and S2**) were rendered by PyMOL and Adobe Photoshop 2023.

## Supporting information

Movie S1 (PC1 motion)

Movie S2 (PC2 motion)

## SUPPLEMENTAL INFORMATION

### ACKNOWLEDGEMENTS

We thank Dr. Thomas Klose in the Purdue Cryo-EM Facility for technical assistance and Dr. Vsevolod V. Gurevich in the Department of Pharmacology from Vanderbilt University for generous gifts of Arr2 and Arr3 plasmids. This work was supported by the Center for Electron Microscopy (iCEM) at Indiana University School of Medicine. Support from Purdue Institute for Cancer Research Phase I and II Pilot Awards is gratefully acknowledged, P30CA023168. Funding was provided by:

National Institutes of Health grant AI161880 (TMH)

National Institutes of Health grant CA254402 (JJGT, TMH)

National Institutes of Health grant CA221289 (JJGT)

National Institutes of Health grant CA023168 (JJGT)

National Institutes of Health grant HL071818 (JJGT)

National Institutes of Health grant P30CA023168 (JJGT)

National Institutes of Health grant GM117372 (AAK)

Walther Cancer Foundation (JJGT)

National Institutes of Health grant F32 GM137505 (CTS)

Robertson Foundation/Cancer Research Institute Irvington Postdoctoral Fellowship (MG)

VILLUM FONDEN research grant 00025326 (MG)

### AUTHOR CONTRIBUTIONS

Q.C., J.J.G.T. and T.M.H. conceptualized the study. C.T.S. and M.G. expressed and purified ACKR3 variants. Q.C. produced and purified arrestins, GRK2, and GRK5. C.T.S., M.G. and Q.C. performed pulldown assays. Q.C. labelled arrestins and performed the fluorescence measurements. S.M., P.A., Q.C., and A.A.K. selected the Fabs. Q.C. and J.J.G.T. collected data and performed structure determinations of all cryo-EM structures. X.-Q.Y. performed the PCA analysis. Q.C. wrote the original draft and all authors further edited the manuscript. Q.C., J.J.G.T. and T.M.H contributed funding.

